# DepEst: an R package of important dependency estimators for gene network inference algorithms

**DOI:** 10.1101/102871

**Authors:** Gökmen Altay, Zeyneb Kurt, Nejla Altay, Nizamettin Aydin

## Abstract

Gene network inference algorithms (GNI) are popular in bioinformatics area. In almost all GNI algorithms, the main process is to estimate the dependency (association) scores among the genes of the dataset.

We present a bioinformatics tool, DepEst (Dependency Estimators), which is a powerful and flexible R package that includes 11 important dependency score estimators that can be used in almost all GNI Algorithms. DepEst is the first bioinformatics package that includes such a large number of estimators that runs both in parallel and serial.

DepEst is currently available at https://github.com/altayg/Depest. Package access link, instructions, various workflows and example data sets are provided in the supplementary file.

## 1. INTRODUCTION

Gene network inference algorithms are very popular in bioinformatics research area. Those algorithms provide us to explore the vast amount of the interactions among the molecules in the cell. Application areas of GNI algorithms are very wide; for example discovering a genome-wide genes and gene-products interactions, determining the main target of a drug in pharmacological studies and so on.

The most crucial process of the GNI is to estimate the interaction scores among the cell molecules such as genes or proteins. It is a challenging process because of the current very large-scale biological datasets and the noise caused by the experimental and computational processes. If this step is not correctly fulfilled then the rest of inference process becomes erroneous no matter which GNI algorithm is used [1,2,3,4].

In [5], almost all the important estimators in literature are compared and concluded the outperforming estimators. Here we present an R package that includes 11 outperforming dependency score estimators determined in that comparison study. This package can be used in all the GNI algorithms to replace the estimator part with respect to the estimator performances of [5,6,7], so that it may enhance the overall GNI performance. Our package also saves time since it may also run in parallel if needed. Nowadays the datasets get larger therefore this property can also be very helpful.

## 2. METHODS

DepEst is a comprehensive tool that can be used to compare and evaluate dependency score estimators. Furthermore the performance of different GNI algorithms can be assessed through the package for each dependency score estimator. The R package contains 11 important dependency estimators [5] that works in parallel or in serial programming which are as follows:

- Pearson Correlation Coefficient (PCC)
- Spearman Correlation Coefficient (SCC)
- Pearson Based Gaussian Estimator (PBG)
- Spearman Based Gaussian Estimator (SBG)
- Nth Order Partial Pearson Correlation Coefficient (PPCN)
- Heller Heller Gorfine Estimator (HHG)
- Miller-Madow Mutual Information Estimator (MM)
- Chao-Shen Mutual Information Estimator (CS)
- B-spline Mutual Information Estimator (BS)
- Kernel Density Estimator (KDE)
- K Nearest Neighborhood Estimator (KNN)

The R package also includes auxiliary functions for equal frequency discretization, equal wideness discretization, elimination of non-significant interactions, copula transformation and cluster installation for parallel computing. The PCC, SCC, PPCN and HHG methods of the package obtain association scores using correlation-based approach. The rest of methods use mutual information (MI) based approach [5].

DepEst provides obtain.mim function to compute the mutual information matrix of the genes (random variables) by using the selected estimator. The following command would be an example of a call to obtain.mim to obtain a mutual information matrix using Miller Madow estimator:

`mim <- obtain.mim(expdataset, estimator="miller.madow", cop.transform= TRUE, disc.method="eq.freq", parallel=FALSE)`

The required inputs of the function are the gene expression dataset (expdataset), the name of the estimator, and required arguments of the specified estimator. Optional input parameters that can be used with obtain.mim function are cop.transform, num.of.bins, disc.method, bandwidth, spline.order, num.of.neighbours and parallel. Here, the first parameter controls the preprocessing step that provides more stable estimations by setting the parameter as TRUE. The num.of.bins is the number of bins to be used for discretization process. The disc.method is the name of the discretization method for the entropy based MI estimators (MM and CS). The bandwidth parameter determines the amount of smoothness of the estimated density, when the KDE is used. Also it strongly affects the accuracy of the estimator. The spline.order is the order of the spline to be used for the B spline estimator. It denotes the number of bins each sample belongs in. The num.of.neighbours specifies the neighborhood for the KNN Estimator. The parallel parameter denotes that whether parallel or serial version of the specified estimator is used. This is a key feature of our package that decreases computation time significantly.

Since DepEst is not a gene network inference algorithm, users who want to visualize a network should first infer a gene network using one of gene network inference methods and then a visualization package such as igraph [8], Cytoscape [9] or ReDer [10] can be used to visualize the gene network. An example using gene network inference method C3NET [1] is provided at below.

`library(c3net)`

`net <- c3(mim)`

`netplot(net)`

The comparison and evaluation of the dependency score estimators was performed by using the GNI algorithms ARACNE [6] and C3NET [1] with a real biological Escherichia coli dataset [11]. This data set has 3091 validations, which can be used to evaluate performance of dependency estimators using a real dataset. Since the true network of the dataset is not complete and only the known interactions are used, Precision metric is used in the comparison.

**Figure 1.**
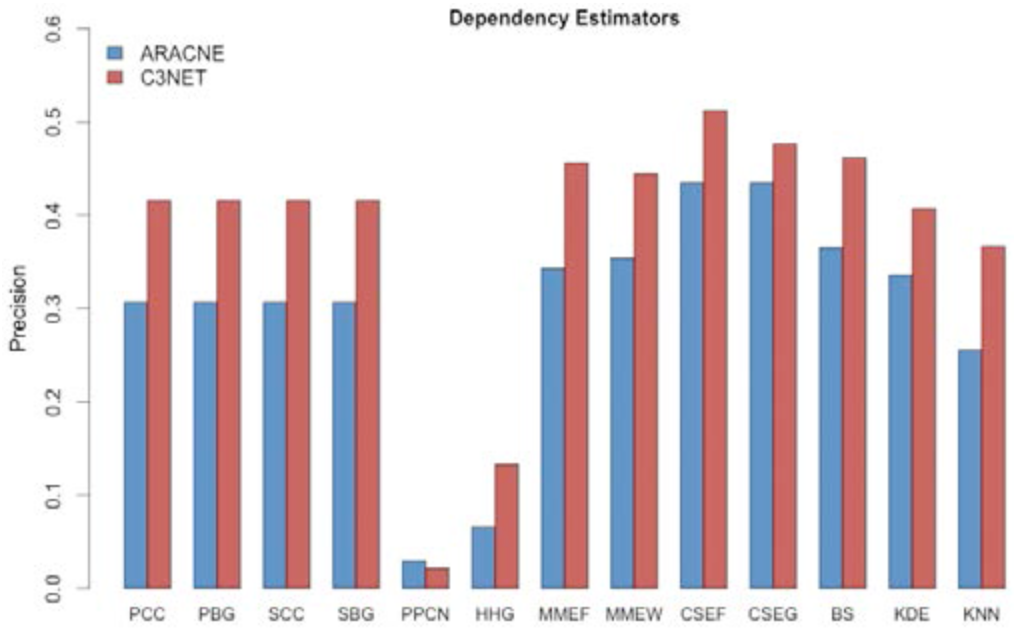
Precision scores of the estimators for the E.coli dataset.

## 3. RESULTS

In this section, an example evaluation of the dependency estimators using DepEst package is shown. Further detailed performance comparison analysis can be seen in [5]. The real biological Escherichia coli dataset involves 524 microarray chips and 1146 genes. For this dataset, barplot of precision values of the estimators according to the two GNI algorithms is given in Figure 1.

As shown in Figure 1, CSEF and CSEW are the most promising estimators for ARACNE (EF for equal frequency and EW for equal width). The BS, MMEF, MMEW and KDE follow the CS closely. PCC, PBG, SCC and SBG are the other estimators that perform close to the former ones. However, PPCN, HHG and KNN do not perform well.

When C3NET is used, CSEF and CSEW are the best performing estimators again. BS, MMEF and MMEW closely follow CS method. Therefore, CS, BS and MM become prominent. For C3NET, all the estimators except PPCN and HHG are observed as promising.

## 4. DISCUSSION

We present DepEst, an R package that provides 11 dependency score estimators for various Gene Network Inference Algorithms. Since R is a popular statistical package among biologists, DepEst is available for all platforms, Windows, Mac OS and Linux, through the R user interface.

DepEst is one of the first bioinformatics package that includes such a large number of association estimators in one single package. Furthermore, there is no package that can be run also in parallel for all of the estimators it includes. Parallel computing feature of DepEst saves time by decreasing computation time significantly that makes DepEst powerful and timely.

## SUPPLEMENTARY MATERIAL

Package instructions, various workflows and example data sets are provided in the supplementary file following the manuscript.

## Supplementary File

DepEst: an R package of important dependency estimators for gene network inference algorithms

Package Version: 1.1

January, 2017

## 1. Introduction

Gene network inference algorithms (GNI) are widely used in Bioinformatics area. In almost all GNI algorithms, the main process is to estimate the association scores among the variables of the dataset. The *DepEst* provides a powerful and flexible framework that includes 11 different dependency score estimators for various GNI Algorithms.

This supplementary file exemplifies how to use the *DepEst* package and express detailed information on the several workflows with example data set. The data set used in this file are available through the *DepEst* R-package.

## 2. Installation of the R package DepEst

*DepEst* requires “R 3.2.x and later” and it depends on “parallel”, “foreach”, “snow”, “doSNOW” and “corpcor” packages that can be installed from the CRAN and Bioconductor libraries. *DepEst* also requires “Rtools” installed in the computer for Windows OS. For the installation of *DepEst*, the user needs to follow some simple installation steps. The software will be uploaded to CRAN upon publication.

### 2.1. Installation on Windows

1. Download and install “Rtools33.exe” (R compatibility: R 3.2.x and later) from https://cran.r-project.org/bin/windows/Rtools Please tick up the "Edit the system PATH" checkbox during the installation. After the installation is completed, please restart your computer (required).
2. To download and install dependent packages *c3net, igraph, minet, parallel, snow, doSNOW* and *corpcor* from CRAN and Bioconductor (execute in R): *> source("http://bioconductor.org/biocLite.R")* `> biocLite("parallel")` `> biocLite("foreach")` `> biocLite("snow")` `> biocLite("doSNOW")` `> biocLite("corpcor")`
3. Execute the installation command for *DepEst* in R *> install.packages("https://github.com/altayg/Depest/raw/master/DepEst_1.1.tar.gz", type="source", repos=NULL)*
4. For the instructions on the usage of *DepEst*, please check the user manual *DepEst-manual.pdf.* `> help(package=DepEst)`
5. To load the library: `> library(DepEst)`

### 2.2. Installation on Mac or Linux

1. To download and install dependent packages c3net, igraph, minet, parallel, snow, doSNOW and corpcor from CRAN and Bioconductor (execute in R): *> source("http://bioconductor.org/biocLite.R")* `> biocLite("parallel")` `> biocLite("foreach")` `> biocLite("snow")` `> biocLite("doSNOW")` `> biocLite("corpcor")`
2. Execute the installation command for DepEst in R *> install.packages("https://github.com/altayg/Depest/raw/master/DepEst_1.1.tar.gz", type="source", repos=NULL)*
3. For the instructions on the usage of DepEst, please check the user manual DepEst-manual.pdf. `> help(package=DepEst)`
4. To load the library: `> library(DepEst)`

## 3. General guidelines for using DepEst

The R package *DepEst* contains 11 different association estimators that works in parallel or in series programming which are as follows:

- Pearson Correlation Coefficient (PCC)
- Spearman Correlation Coefficient (SCC)
- Pearson Based Gaussian Estimator (PBG)
- Spearman Based Gaussian Estimator (SBG)
- Nth Order Partial Pearson Correlation Coefficient (PPCN)
- Miller-Madow Mutual Information Estimator (MM)
- Chao-Shen Mutual Information Estimator (CS)
- B-spline Mutual Information Estimator (BS)
- Heller Heller Gorfine Estimator (HHG)
- Kernel Density Estimator (KDE)
- K Nearest Neighborhood Estimator (KNN)

The R package also includes auxiliary functions for equal frequency discretization, equal wideness discretization, elimination of non-significant interactions, copula transformation and cluster installation for parallel computing. The PCC, SCC, PPCN and HHG methods of the package obtain association scores using correlation-based approach. The rest of methods use mutual information (MI) based approach (Kurt et al., 2014).

Since R is an interpreted language, the core functions of R give fast results. Contrary to this, user-defined functions work slow and the results are not fast as the core functions. Therefore, we implemented BS, CS, KDE, HHG, KNN and MM methods in C++ using parallel programming to faster the computation time. For the PCC, SCC, PBGE, SBGE and PPCCn methods, we used the core functions of R.

DepEst provides `obtain.mim` function to compute the mutual information matrix of the genes (random variables) by using the selected estimator. The required inputs of the function are the gene expression dataset, the name of the estimator, and required arguments of the specified estimator. Users may also prefer to use directly the function of the estimator rather than the main function of the package. The details are described in the package manual and in the tutorial section of this document.

`> obtain.mim(dataset, estimator, arguments)`

Here, `dataset` is the gene expression dataset whose mutual information (MI) scores will be obtained by using the selected estimator. The second parameter, `estimator`, is the name of the estimator. The default value for the estimator is "scc". The abbreviations of the selected estimator that must be used with the estimator parameter are:

| Estimator | Abbreviation |
| --- | --- |
| Pearson Correlation Coefficient | pcc |
| Spearman Correlation Coefficient | scc |
| Pearson-Based Gaussian Estimator | pbg |
| Spearman-Based Gaussian Estimator | sbg |
| n-th Order Pearson Correlation Coefficient | ppcn |
| Miller-Madow Estimator | miller.madow |
| Chao-Shen Estimator | chao.shen |
| B-Spline Estimator | b.spline |
| Heller Heller Gorfine Estimator | hhg |
| Kernel Density Estimator | kde |
| K-Nearest Neighbourhood Estimator | knn |

Optional input parameters that can be used with the `obtain.mim` function are:

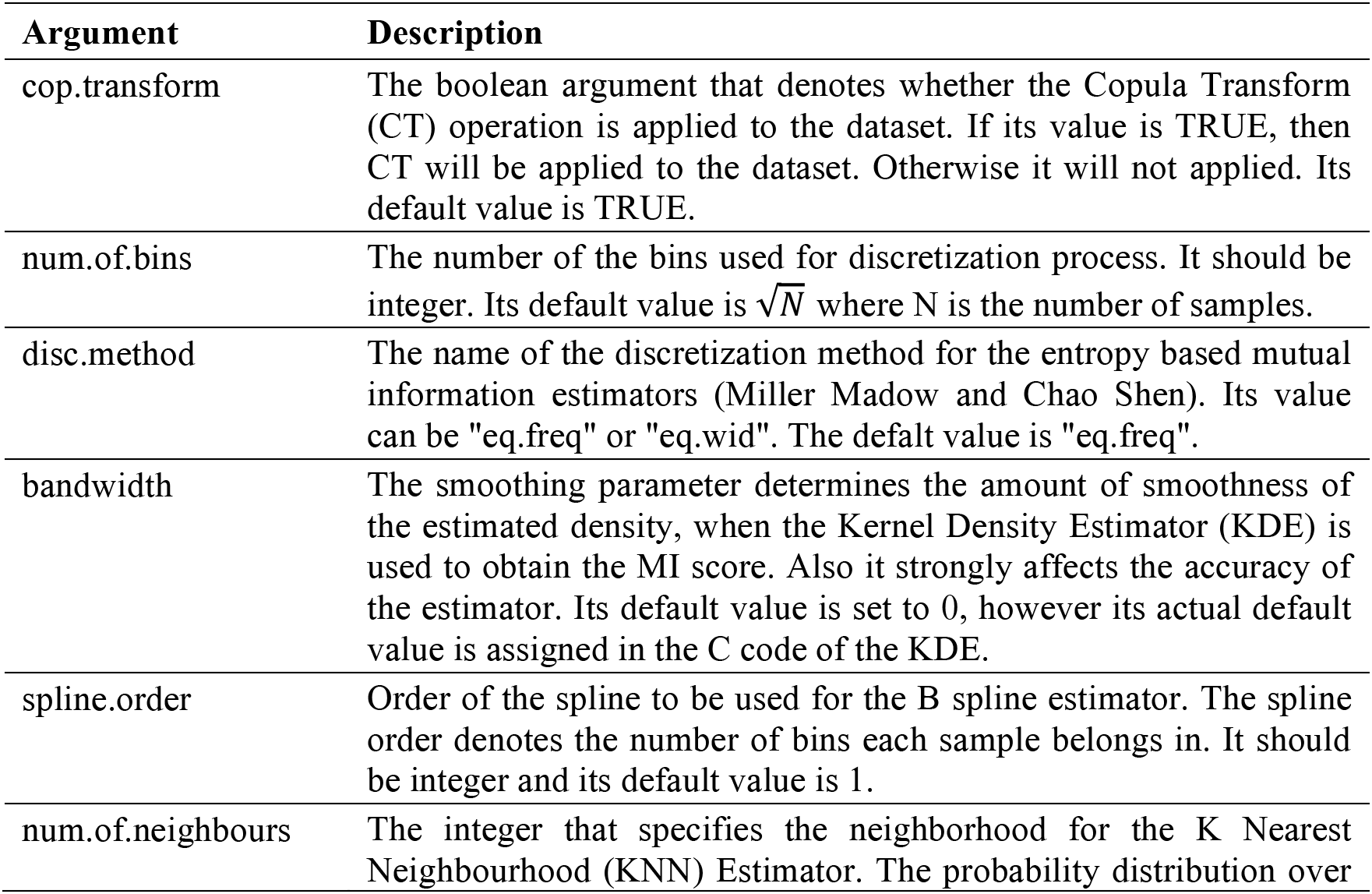

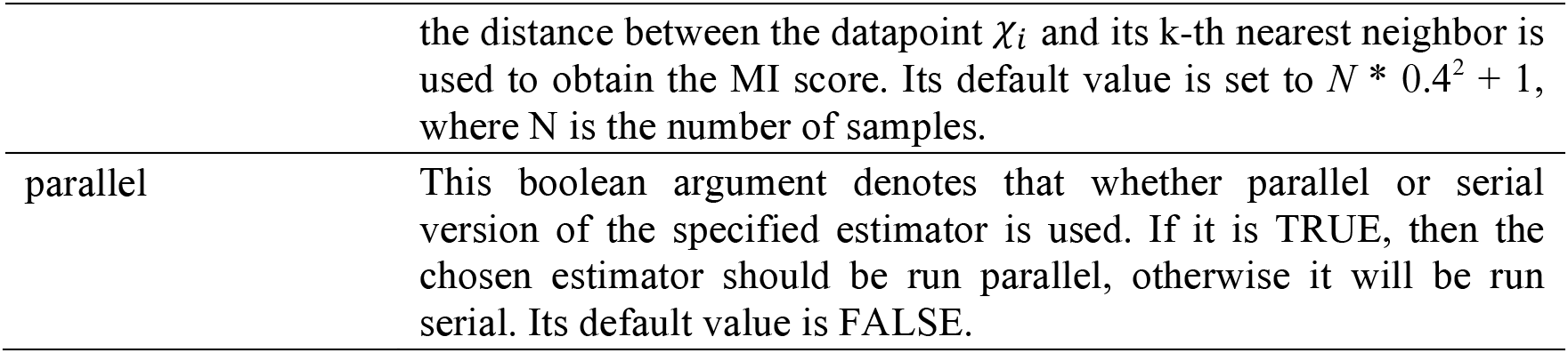

To use the parallel version of the selected estimator, users should first run the `setupCls` function to set the clusters.

`> setupCls(use.cores, cls.list)`

Here, `use.cores` is the number of the nodes (cores) in a work station to be used for running parallel codes. Its default value is the core number of user’s computer minus 1. `cls.list` is the list of the clusters to be used for running parallel codes. If it is not specified, `setupCls` function assumes that, single machine is used for parallel codes and default core numbers denoted above (argument `use.cores`) is used. Parallel, snow, and doSNOW packages are used to contruct the cluster. That cluster will be used to run the parallel implemented versions of the estimators.

Finally, `obtain.mim` function returns the mutual information matrix (MIM) by using the selected method and method specific parameters. MIM is a square matrix that contains the interaction scores among the gene pairs and whose (i, j) element is the mutual information score between the *Gene*_*i*_ and *Gene*_*j*_.

The detailed information about the functions and parameters are described in the manual file, “DepEst-manual.pdf”, which can be accessed through the R package.

## 4. Data structure

Load the example data set contained in the *DepEst* package:

`> ### Load the example data set`

`> data(expdataset)`

The object `expdataset` is the gene expression dataset that involves each gene (random variable) in each row and contains each sample in each column. The dimensions of the example data set is 100x100.

`> ### The dimension of the example data set`

`> dim(expdataset)`

`[1] 100 100`

## 5. Tutorial

### 5.1. Usage of Association Estimators

In this part, we show step by step, how to obtain MI matrices using the selected estimator. The commands below are essential for all of the estimators to load the *DepEst* package and example data set.

`> library(DepEst)`

`> data(expdataset)`

#### 5.1.1. Pearson Correlation Coefficient Estimator

PCC Estimator can only obtain the linear correlation scores among the random variables (genes). Usage of Pearson Correlation Coefficient Estimator with `obtain.mim` function:

`> data(expdataset)`

`> mim <- obtain.mim(expdataset, estimator="pcc")`

Alternative usage of Pearson Correlation Coefficient Estimator with `pearson` function:

`> data(expdataset)`

`> mim <- pearson(expdataset)`

Pre-processing can be enabled by setting `cop.transform` parameter as TRUE in both functions:

`> data(expdataset)`

`> mim <- obtain.mim(expdataset, estimator="pcc", cop.transform=TRUE)`

or

`> data(expdataset)`

`> mim <- pearson(expdataset, cop.transform=TRUE)`

To run parallel version of Pearson Correlation Coefficient Estimator, clusters should be first setup. The detailed information about the usage of `setupCls` function can be found in the package manual file.

`> data(expdataset)`

`> cls.pr <- setupCls(use.cores = 2)`

`> mim <- obtain.mim(expdataset, estimator="pcc", parallel=TRUE)`

or

`> data(expdataset)`

`> cls.pr <- setupCls(use.cores = 2)`

`> mim <- pearson(expdataset, parallel=TRUE)`

The commands above demonstrates the usage of Pearson Correlation Coefficient Estimator. The execution of the commands returns the MI matrix and assign it to `mim` object.

#### 5.1.2. Spearman Correlation Coefficient Estimator

SCC Estimator can only obtain the monotone correlation scores among the random variables (genes). It could also find the correlation score of the non linear relationships. Usage of Spearman Correlation Coefficient Estimator with `obtain.mim` function:

`> data(expdataset)`

`> mim <- obtain.mim(expdataset, estimator="scc")`

Alternative usage of Sperman Correlation Coefficient Estimator with `spearman` function:

`> data(expdataset)`

`> mim <- spearman(expdataset)`

Pre-processing can be enabled by setting `cop.transform` parameter as TRUE in both functions:

`> data(expdataset)`

`> mim <- obtain.mim(expdataset, estimator="scc", cop.transform=TRUE)`

or

`> data(expdataset)`

`> mim <- spearman(expdataset, cop.transform=TRUE)`

To run parallel version of Spearman Correlation Coefficient Estimator:

`> data(expdataset)`

`> cls.pr <- setupCls(use.cores = 2)`

`> mim <- obtain.mim(expdataset, estimator="scc", parallel=TRUE)`

or

`> data(expdataset)`

`> mim <- spearman(expdataset, parallel=TRUE)`

The commands above demonstrates the usage of Spearman Correlation Coefficient Estimator. The execution of the commands returns the MI matrix and assign it to `mim` object.

#### 5.1.3. Pearson-Based Gaussian Estimator

PBG is based on PCC method and uses the correlation scores to calculate the MI scores. It obtains the MI scores between two random variables by assuming that the joint probability among these variables is distributed according to the Gaussian (normal) function. Usage of Pearson-Based Gaussian Estimator with `obtain.mim` function:

`> data(expdataset)`

`> mim <- obtain.mim(expdataset, estimator="pbg")`

Alternative usage of Pearson-Based Gaussian Estimator with `pearson.based.gauss` function:

`> data(expdataset)`

`> mim <- pearson.based.gauss(expdataset)`

Pre-processing can be enabled by setting `cop.transform` parameter as TRUE in both functions:

`> data(expdataset)`

`> mim <- obtain.mim(expdataset, estimator="pbg", cop.transform=TRUE)`

or

`> data(expdataset)`

`> mim <- pearson.based.gauss(expdataset, cop.transform=TRUE)`

To run parallel version of Pearson-Based Gaussian Estimator:

`> data(expdataset)`

`> cls.pr <- setupCls(use.cores = 2)`

`> mim <- obtain.mim(expdataset, estimator="pbg", parallel=TRUE)`

or

`> data(expdataset)`

`> mim <- pearson.based.gauss(expdataset, parallel=TRUE)`

The commands above demonstrates the usage of Pearson-Based Gaussian Estimator. The execution of the commands returns the MI matrix and assign it to `mim` object.

#### 5.1.4. Spearman-Based Gaussian Estimator

SBG is based on SCC method and uses the correlation scores to calculate the MI scores. It obtains the MI scores between two random variables by assuming that the joint probability among these variables is distributed according to the Gaussian (normal) function. Usage of Spearman-Based Gaussian Estimator with `obtain.mim` function:

`> data(expdataset)`

`> mim <- obtain.mim(expdataset, estimator="sbg")`

Alternative usage of Spearman-Based Gaussian Estimator with `spearman.based.gauss` function:

`> data(expdataset)`

`> mim <- spearman.based.gauss(expdataset)`

Pre-processing can be enabled by setting `cop.transform` parameter as TRUE in both functions:

`> data(expdataset)`

`> mim <- obtain.mim(expdataset, estimator="sbg", cop.transform=TRUE)`

or

`> data(expdataset)`

`> mim <- spearman.based.gauss(expdataset, cop.transform=TRUE)`

To run parallel version of Spearman-Based Gaussian Estimator:

`> data(expdataset)`

`> cls.pr <- setupCls(use.cores = 2)`

`> mim <- obtain.mim(expdataset, estimator="sbg", parallel=TRUE)`

or

`> data(expdataset)`

`> mim <- spearman.based.gauss(expdataset, parallel=TRUE)`

The commands above demonstrates the usage of Spearman-Based Gaussian Estimator. The execution of the commands returns the MI matrix and assign it to `mim` object.

#### 5.1.5. n-th Order Pearson Correlation Coefficient Estimator

PPCN not only obtains the correlation scores among the genes, it can also eliminate the indirected edges in the gene network. It is alternatively called as Graphical Gaussian Modeling (GGM) method. It computes the inverse of the 0-th order Pearson Correlation matrix and normalizing this inverse matrix to have diagonals -1. Usage of n-th Order Pearson Correlation Coefficient Estimator with `obtain.mim` function:

`> data(expdataset)`

`> mim <- obtain.mim(expdataset, estimator="ppcn")`

Alternative usage of n-th Order Pearson Correlation Coefficient Estimator with `ppcn` function:

`> data(expdataset)`

`> mim <- ppcn(expdataset)`

Pre-processing can be enabled by setting `cop.transform` parameter as TRUE in both functions:

`> data(expdataset)`

`> mim <- obtain.mim(expdataset, estimator="ppcn", cop.transform=TRUE)`

or

`> data(expdataset)`

`> mim <- ppcn(expdataset, cop.transform=TRUE)`

To run parallel version of n-th Order Pearson Correlation Coefficient Estimator:

`> data(expdataset)`

`> cls.pr <- setupCls(use.cores = 2)`

`> mim <- obtain.mim(expdataset, estimator="ppcn", parallel=TRUE)`

or

`> data(expdataset)`

`> mim <- ppcn(expdataset, parallel=TRUE)`

The commands above demonstrates the usage of n-th Order Pearson Correlation Coefficient Estimator. Ppcn returns the N-th order partial correlation matrix and assign it to `mim` object.

#### 5.1.6. Miller-Madow Estimator

Miller-Madow estimator is a bias-corrected version of maximum likelihood mutual information estimator. The required arguments for this estimator are `num.of.bins` and `disc.method. num.of.bins` is the number of the bins used for discretization process. It should be integer. Its default value is 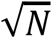 where N is the number of samples. `disc.method` is the name of the discretization method. Its value can be "eq.freq" or "eq.wid". The defalt value is "eq.freq".

Usage of Miller-Madow Estimator with `obtain.mim` function:

`> data(expdataset)`

`> mim <- obtain.mim(expdataset, estimator="miller.madow", disc.method="eq.freq")`

Alternative usage of Miller-Madow Estimator with `miller.madowR` function:

`> data(expdataset)`

`> mim <- miller.madowR(expdataset, disc.method="eq.freq")`

Pre-processing can be enabled by setting `cop.transform` parameter as TRUE in both functions:

`> data(expdataset)`

`> mim <- obtain.mim(expdataset, estimator="miller.madow", disc.method="eq.freq", cop.transform=TRUE)`

or

`> data(expdataset)`

`> mim <- miller.madowR(expdataset, disc.method="eq.freq", cop.transform=TRUE)`

To run parallel version of Miller-Madow Estimator:

`> data(expdataset)`

`> cls.pr <- setupCls(use.cores = 2)`

`> mim <- obtain.mim(expdataset, estimator="miller.madow", disc.method="eq.freq", parallel=TRUE)`

or

`> data(expdataset)`

`> mim <- miller.madowParR(expdataset, disc.method="eq.freq")`

`miller.madowR` function returns the MI matrix by using serial version of Miller-Madow Estimator. `miller.madowParR` function should be run to use parallel version.

#### 5.1.7. Chao-Shen Estimator

Chao-Shen estimator combines Horvitz Thompson estimator and Good Turing correction of the Maximum Likelihood estimators. It is proposed by Chao and Shen. The required arguments for this estimator are `num.of.bins` and `disc.method`. `num.of.bins` is the number of the bins used for discretization process. It should be integer. Its default value is 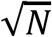 where N is the number of samples. `disc.method` is the name of the discretization method. Its value can be "eq.freq" or "eq.wid". The defalt value is "eq.freq".

Usage of Chao-Shen Estimator with `obtain.mim` function:

`> data(expdataset)`

`> mim <- obtain.mim(expdataset, estimator="chao.shen", disc.method="eq.freq")`

Alternative usage of Chao-Shen Estimator with `chao.shenR` function:

`> data(expdataset)`

`> mim <- chao.shenR(expdataset, disc.method="eq.freq")`

Pre-processing can be enabled by setting `cop.transform` parameter as TRUE in both functions:

`> data(expdataset)`

`> mim <- obtain.mim(expdataset, estimator="chao.shen", disc.method="eq.freq", cop.transform=TRUE)`

or

`> data(expdataset)`

`> mim <- chao.shenR(expdataset, disc.method="eq.freq", cop.transform=TRUE)`

To run parallel version of Chao-Shen Estimator:

`> data(expdataset)`

`> cls.pr <- setupCls(use.cores = 2)`

`> mim <- obtain.mim(expdataset, estimator="chao.shen", disc.method="eq.freq", parallel=TRUE)`

or

`> data(expdataset)`

`> mim <- chao.shenParR(expdataset, disc.method="eq.freq")`

`chao.shenR` function returns the MI matrix by using serial version of Chao-Shen Estimator. `chao.shenParR` function should be run to use parallel version.

#### 5.1.8. B-Spline Estimator

The spline order denotes the number of bins each sample belongs in. Hence, B-spline provides soft binning operation rather than hard binning, unlike the other mutual information based methods, which requires binning operation.

B-Spline estimator combines Horvitz Thompson estimator and Good Turing correction of the Maximum Likelihood estimators. It is proposed by Chao and Shen. The required arguments for this estimator are `num.of.bins` and `spline.order. num.of.bins` is the number of the bins used for discretization process. It should be integer. Its default value is 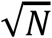 where N is the number of samples. `spline.order` is the order of the spline to be used for the B spline estimator. The spline order denotes the number of bins each sample belongs in. It should be integer and its default value is 1.

Usage of B-Spline Estimator with `obtain.mim` function:

`> data(expdataset)`

`> mim <- obtain.mim(expdataset, estimator="b.spline", spline.order=1)`

Alternative usage of B-Spline Estimator with `chao.shenR` function:

`> data(expdataset)`

`> mim <- b.splineR(expdataset, spline.order=1)`

Pre-processing can be enabled by setting `cop.transform` parameter as TRUE in both functions:

`> data(expdataset)`

`> mim <- obtain.mim(expdataset, estimator="b.spline", spline.order=1, cop.transform=TRUE)`

or

`> data(expdataset)`

`> mim <- b.splineR(expdataset, spline.order=1, cop.transform=TRUE)`

To run parallel version of B-Spline Estimator:

`> data(expdataset)`

`> cls.pr <- setupCls(use.cores = 2)`

`> mim <- obtain.mim(expdataset, estimator="b.spline", spline.order=1, parallel=TRUE)`

or

`> data(expdataset)`

`> mim <- b.splineParR(expdataset, spline.order=1)`

`b.splineR` function returns the MI matrix by using serial version of B-Spline Estimator. `b.splineParR` function should be run to use parallel version.

#### 5.1.9. Heller Heller Gorfine Estimator

HHG is a metric that measures the independency among the random variables. Usage of Heller Heller Gorfine Estimator with `obtain.mim` function:

`> data(expdataset)`

`> mim <- obtain.mim(expdataset, estimator="hhg")`

Alternative usage of Heller Heller Gorfine Estimator with `chao.shenR` function:

`> data(expdataset)`

`> mim <- hhgR(expdataset)`

Pre-processing can be enabled by setting `cop.transform` parameter as TRUE in both functions:

`> data(expdataset)`

`> mim <- obtain.mim(expdataset, estimator="hhg", cop.transform=TRUE)`

or

`> data(expdataset)`

`> mim <- hhgR(expdataset, cop.transform=TRUE)`

To run parallel version of Heller Heller Gorfine Estimator:

`> data(expdataset)`

`> cls.pr <- setupCls(use.cores = 2)`

`> mim <- obtain.mim(expdataset, estimator="hhg", parallel=TRUE)`

or

`> data(expdataset)`

`> mim <- hhgParR(expdataset)`

`hhgR` function returns the MI matrix by using serial version of Heller Heller Gorfine Estimator. `hhgParR` function should be run to use parallel version.

#### 5.1.10. Kernel Density Estimator

Gaussian function is used as kernel in KDE. Hence the bandwidth parameter, h, is the standard deviation of the Gaussian function. The required argument for this estimator is `bandwidth. bandwidth` or in other words the smoothing parameter determines the amount of smoothness of the estimated density. Also it strongly affects the accuracy of the estimator. Usage of Kernel Density Estimator with `obtain.mim` function:

`> data(expdataset)`

`> mim <- obtain.mim(expdataset, estimator="kde", bandwidth=0)`

Alternative usage of Kernel Density Estimator with `kdeR` function:

`> data(expdataset)`

`> mim <- kdeR(expdataset, bandwidth=0)`

Pre-processing can be enabled by setting `cop.transform` parameter as TRUE in both functions:

`> data(expdataset)`

`> mim <- obtain.mim(expdataset, estimator="kde", bandwidth=0, cop.transform=TRUE)`

or

`> data(expdataset)`

`> mim <- kdeR(expdataset, bandwidth=0, cop.transform=TRUE)`

To run parallel version of Kernel Density Estimator:

`> data(expdataset)`

`> cls.pr <- setupCls(use.cores = 2)`

`> mim <- obtain.mim(expdataset, estimator="kde", bandwidth=0, parallel=TRUE)`

or

`> data(expdataset)`

`> mim <- kdeParR(expdataset, bandwidth=0)`

`kdeR` function returns the MI matrix by using serial version of Kernel Density Estimator. kdeParR function should be run to use parallel version.

#### 5.1.11. K Nearest Neighborhood Estimator

KNN Estimator directly obtains the mutual information (MI) score between two random variables. It does not require the entropy calculation. The probability distribution over the distance between the datapoint X_i_ and its k-th nearest neighbor is used to obtain the MI score. The required argument for this estimator is `num.of.neighbours. num.of.neighbours` is the integer that specifies the neighborhood. Usage of K Nearest Neighborhood Estimator with `obtain.mim` function:

`> data(expdataset)`

`> mim <- obtain.mim(expdataset, estimator="knn", num.of.neighbours=0)`

Alternative usage of K Nearest Neighborhood Estimator with `knn.direct.miR` function:

`> data(expdataset)`

`> k <- 0`

`> mim <- knn.direct.miR(expdataset, k)`

Pre-processing can be enabled by setting `cop.transform` parameter as TRUE in both functions:

`> data(expdataset)`

`> mim <- obtain.mim(expdataset, estimator="knn", num.of.neighbours=0, cop.transform=TRUE)`

or

`> data(expdataset)`

`> mim <- knn.direct.miR(expdataset, cop.transform=TRUE, 0)`

To run parallel version of K Nearest Neighborhood Estimator:

`> data(expdataset)`

`> cls.pr <- setupCls(use.cores = 2)`

`> mim <- obtain.mim(expdataset, estimator="knn", num.of.neighbours=0, parallel=TRUE)`

or

`> data(expdataset)`

`> mim <- knn.direct.miParR(expdataset, cop.transform=TRUE, 0)`

`knn.direct.miR` function returns the MI matrix by using serial version of K Nearest Neighborhood Estimator.

`knn.direct.miParR` function should be run to use parallel version.

### 5.2. Elimination of non-significant edges

In *DepEst* package, we provide `eliminate.nonsignificant.edges` function that takes a MI matrix as input and eliminates its nonsignificant edges by using a simple elimination procedure. Users can determine a particular *Ic* value to eliminate the edges in the MIM by using this function. If *Ic* is not specified, then the value of parameter method determines the elimination procedure.

You can use the output MI matrix obtained from the estimator functions above.

`> data(expdataset)`

`> mim <- eliminate.nonsignificant.edges(mim)`

### 5.3. Comparing the performance scores of GNI algorithms

In this part, we demonstrate how to compare the performance of different GNI algorithms with the selected estimator using *DepEst* package. *minet* and *c3net* packages are required for the GNI algorithms used in this example. To install *c3net* and *minet* packages (execute in R):

`> install.packages("c3net")`

*> source("http://bioconductor.org/biocLite.R")*

`> biocLite("minet")`

In this example parallel version of B-spline Mutual Information Estimator was used to obtain the MI matrix. The commands used are as follows:

`> library(DepEst)`

`> data(expdataset)`

`> data(truenetwork)`

`> mim <- obtain.mim(expdataset, estimator="b.spline", spline.order=1, cop.transform=TRUE, parallel=TRUE)`

`> mim <- eliminate.nonsignificant.edges(mim)`

`> relnet <- mim # Gene network is inferred using Relevance Network`

`> output.relnet <- checknet(relnet, truenetwork) # Get performance scores`

`> eps <- 0`

`> aracne.net <- aracne(mim, eps) # Gene network is inferred using Aracne`

`> output.aracne <- checknet(aracne.net, truenetwork)`

`> c3net <- c3(mim) # Gene network is inferred using C3NET`

`> output.c3net <- checknet(c3net, truenetwork)`

As shown in Figure 1, we obtained three different gene network using Relevance Network, Aracne and C3NET algorithms, respectively. Then we get the performance scores of each GNI algorithm using checknet function. Checknet function computes performance scores by comparing the output GNI networks with the true network.

**Fig 1.**
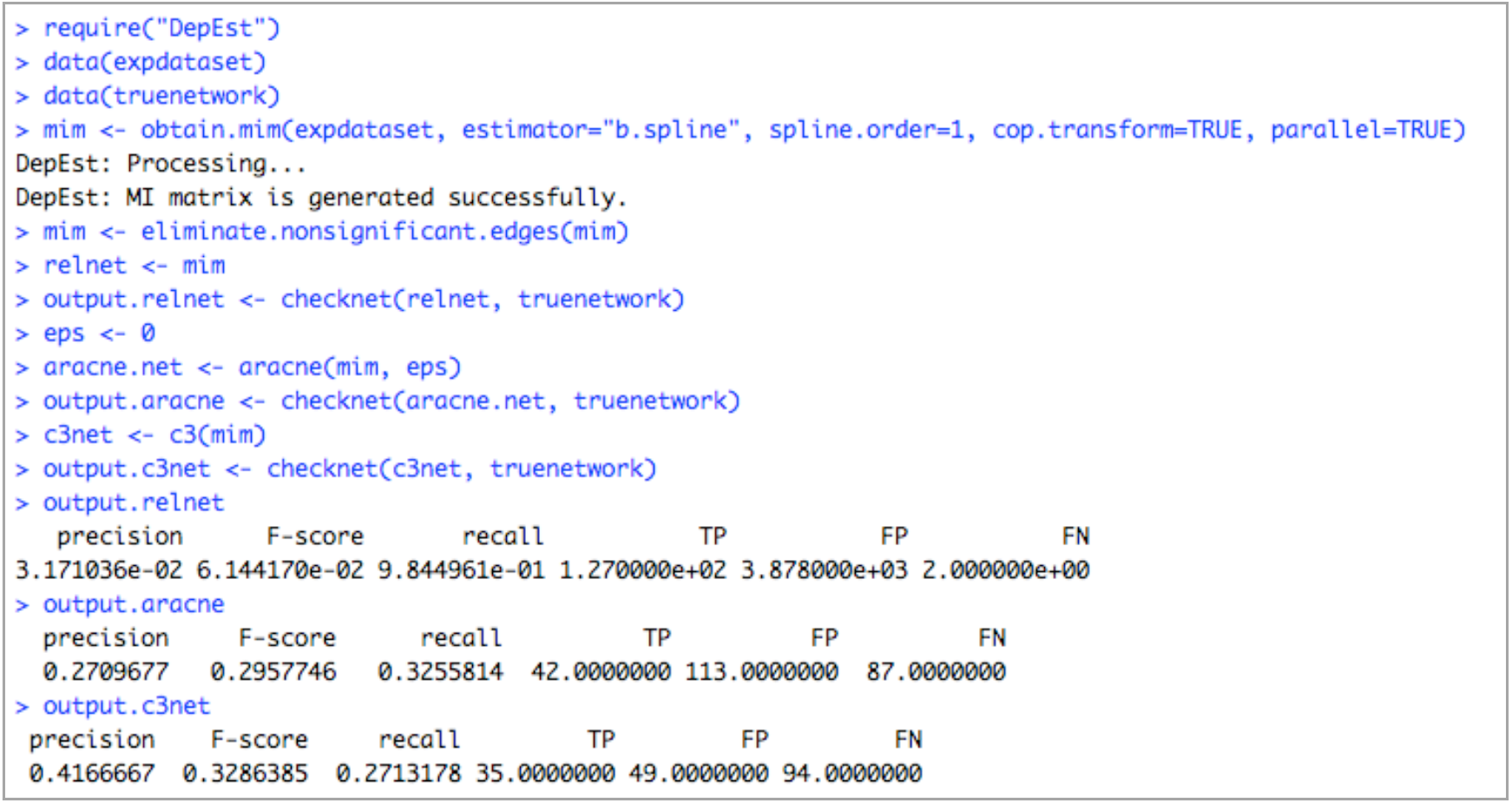
The performance scores of the GNI algorithms

